# Epigenetic Silencing of Carotid Body TRPM7 Attenuates Hypertension in Obese Mice

**DOI:** 10.64898/2026.03.05.709322

**Authors:** Mi-Kyung Shin, Arijit Roy, Omkar Padel, Samhita Gudapati, James SK Sham, Wan-Yee Tang, Vsevolod Y. Polotsky

## Abstract

Obesity is the most common cause of hypertension. We have previously shown that high levels of circulating leptin in diet-induced obese (DIO) mice induced hypertension by increasing expression of Transient Receptor Potential Melastatin-subfamily member 7 (TRPM7) in the carotid bodies (CB). In addition, we demonstrated in rat PC12 cells that leptin increases *Trpm7* gene expression by inducing CpG site-specific demethylation within the 5’ regulatory region containing a signal transducer and activator of transcription 3 (STAT3) binding site. This leptin-induced *Trpm7* upregulation was prevented by inhibition of JAK-STAT3 signaling. Based on these findings, we hypothesized that reversing region-specific methylation at the *Trpm7* promoter in the CB could attenuate obesity-associated hypertension. Compared with lean controls, DIO mice exhibited increased *Trpm7* expression and the STAT3- binding site-specific promoter demethylation in the CB. Administration of methylated DNA oligonucleotides targeting the STAT3 binding site attenuated CpG site-specific DNA demethylation and reduced *Trpm7* transcription in the CB of DIO mice. This intervention resulted in decreased carotid sinus nerve activity and reduced arterial blood pressure, especially during the light phase. Our results suggest that targeted modulation of CpG site-specific DNA methylation at the *Trpm7* promoter using DNA oligonucleotide may represent a novel therapeutic strategy for obesity-induced hypertension.

## Introduction

The prevalence of obesity defined as a body mass index (BMI) ≥ 30 kg/m^2^ has reached 42.4% in the adult US population (1, 2). Obesity markedly increases the risk of hypertension (3–5), which causes high cardiovascular morbidity and mortality (3–11). The prevalence of hypertension in normal-weight subjects is 34% rising to 72.9% in obese individuals (5). At least 20% of hypertensive patients adherent to therapy are resistant to the optimal medical regimens with three or more medications (12). In severely obese individuals, the prevalence of resistant hypertension is 5.3-fold higher than in normal weight patients (5).

A key mechanism by which obesity induces hypertension is activation of the sympathetic nervous system (SNS) (4, 10), either directly or *via* comorbid obstructive sleep apnea (OSA) (10, 13–18). We have previously used a mouse model of diet-induced obesity (DIO) and demonstrated that obesity increases SNS activity and induces hypertension through stimulation of the carotid bodies (CBs) (19–21).

The CB transmits afferent output *via* the carotid sinus nerve (CSN) to the medullary centers that activate the SNS (22–24). The CB plays an important role in the development of hypertension (22, 25, 26). CB denervation effectively controls blood pressure in spontaneously hypertensive rats (27), obese rats (28), obese mice (21), and rodents exposed to intermittent hypoxia (29, 30). Unilateral CB resection decreases SNS activity (31) and leads to sustained blood pressure reductions in patients with resistant hypertension (32, 33).

Obese humans (34, 35) and DIO mice (36, 37) exhibit elevated circulating levels of an adipocyte-derived hormone leptin. Binding of leptin to the long-chain leptin receptor isoform (LEPR^b^) activates Janus kinase 2 (JAK2)-dependent phosphorylation of signal transducer and activator of transcription 3 (pSTAT3), which translocates to the nucleus and activates transcription (38, 39). Leptin increases SNS activity and blood pressure in obese humans and rodents in a dose-dependent manner (4, 40–44). We previously reported high expression of LEPR^b^ in the CB (19, 45), which is further increased in DIO mice compared with lean controls (36) . Leptin binding to CB LEPR^b^ increases CSN activity (45) and induces hypertension (19); these effects can be attenuated by the LEPR^b^ antagonist Allo-Aca (46) and by knockdown of the *Lepr*^*b*^ mRNA in the CB (47).

Leptin-LEPR^b^ signaling in the CB increases expression and activity of the transient receptor potential melastatin-subfamily member 7 (TRPM7) (19). LEPR^b^ and TRPM7 colocalize in tyrosine hydroxylase (TH+) type I glomus cells. Genetic and pharmacological knockdown of TRPM7 in the CB attenuates leptin- and obesity-induced hypertension (19, 20). DIO mice exhibit a 3-fold increase in *Trpm7* mRNA levels in CB (36). Using rat adrenal pheochromocytoma cells stably expressing LEPR^b^ (PC12^LEPRb^) as an *in vitro* model of CB glomus cells, we demonstrated that leptin increases *Trpm7* mRNA expression and induces demethylation of the *Trpm7* promoter at the pSTAT3 binding site (48). These findings suggest that leptin signaling epigenetically regulates *Trpm7* transcription, and that targeted epigenetic modulation may represent a novel therapeutic strategy for TRPM7-mediated hypertension in obesity.

Epigenetic mechanisms, including DNA methylation and histone modifications, are increasingly exploited in drug development (49, 50), with several DNA methyltransferase (DNMT) inhibitors and histone deacetylase inhibitors approved by the FDA for cancer treatment (50). Transcription factor (TF) modifiers are also used to treat diseases associated with dysregulated gene expression (51). However, systemic epigenetic therapies often lack specificity. To address this problem, we propose a targeted epigenetic intervention at a leptin-induced differentially demethylated region of the *Trpm7* promoter. Previous studies (52, 53), including our own (54), have shown that methylated DNA oligonucleotides can induce CpG site-specific DNA methylation and reduce gene transcription both *in vitro* and *in vivo*.

In the current study, we examined the effects of methylated DNA oligonucleotides targeting the leptin-induced differentially demethylated region (MO-R1) *versus* control methylated oligonucleotides (MO-CTL) administered to the CB of DIO mice. Specifically, we assessed their effects on region-specific DNA methylation of the *Trpm7* promoter, *Trpm7* gene expression in the CB, CSN activity, and blood pressure.

## Results

DIO mice (n = 5) exhibited increased *Trpm7* mRNA levels in the CB compared with lean mice (n=5) **(Figure 1A)**. *In silico* analysis revealed a cluster of CpG-rich regions within the *Trpm7* promoter **(Figure 1B)** containing putative binding sites for STAT3, AP4, and USF1, suggesting that obesity-driven epigenetic changes may regulate *Trpm7* transcription. DNA methylation changes at promoter regions can interfere with transcription factor binding and suppress gene expression (55, 56).

**Figure 1.**
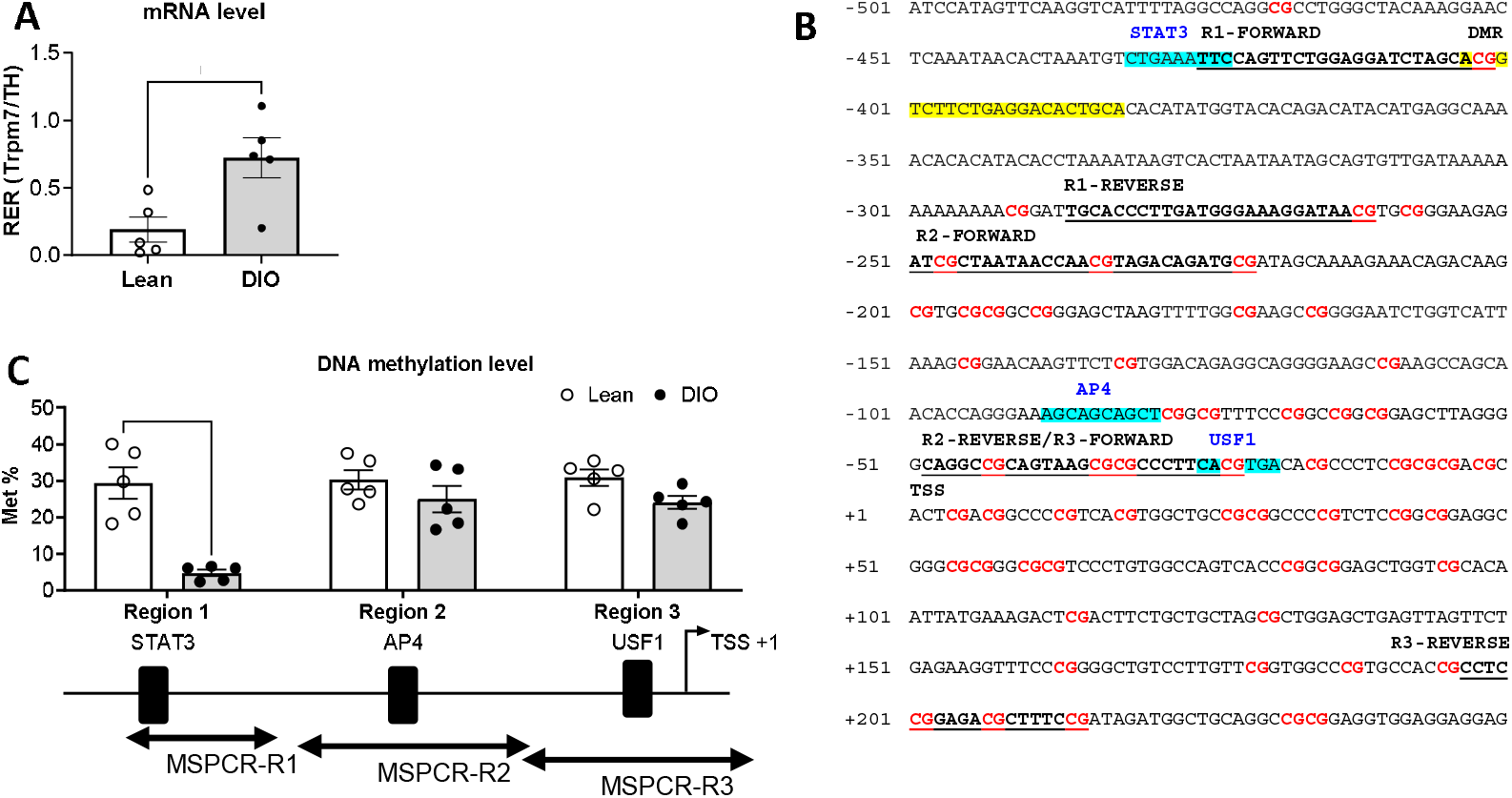
DIO mice exhibit increased Trpm7 mRNA levels and decreased CpG site-specific DNA methylation in CB tissues. **(A)** Comparison of Trpm7 mRNA levels in lean and diet-induced obese (DIO) mice measured by qPCR. **(B)** *In silico* analysis of the mouse Trpm7 promoter showing CpG sites (highlighted in red) and putative transcription factor (TF) binding sites (highlighted in blue). Region-specific DNA methylation of the Trpm7 promoter was analyzed across regions R1, R2, and R3, which encompass STAT3, AP4, and USF1 binding sites, respectively. Primers used for Methylation-Specific PCR (MSPCR) are indicated in bold and underlined. The differentially methylated region (DMR) at R1 is highlighted in yellow; this sequence was used to generate the methylated DNA oligonucleotides for in vivo experiments. **(C)** DNA methylation levels of specific Trpm7 promoter regions assessed by MSPCR. Each dot represents an individual mouse. Data are presented as mean ± SEM.

To identify region-specific DNA methylation changes in DIO mice, we designed primers specific for regions encompassing STAT3, AP4, and USF1. Compared with lean mice, DIO mice exhibited a > 5-fold reduction in DNA methylation at R1 (**Figure 1C**), which encompasses the STAT3 binding site. In contrast, no significant changes were observed at R2 or R3. These findings indicate that CpG site-specific demethylation at R1 contributes to increased Trpm7 transcription in DIO mice.

To establish a causal relationship between CpG site methylation and leptin-induced *Trpm7* transcription, we used methylated DNA oligonucleotides targeting the demethylated CpG sites in PC12^LEPRb^ cells. As previously reported, leptin induces *Trpm7* gene transcription and CpG site-specific demethylation at 5’ regulatory region *via* JAK-STAT signaling (48). In the present study, we generated methylated DNA oligonucleotides (rMO-R1) to target leptin-induced demethylated CpG sites at rat *Trpm7* promoter and a non-targeting methylated control oligonucleotide (rMO-CTL), which is not related to *Trpm*7 but contained methylated cytosine (m5C) residues as the rMO-R1. In the absence of leptin, *Trpm7* mRNA levels were similar in cells treated with rMO-CTL or rMO-R1. Leptin treatment (1 ng/ml for 2 days) increased *Trpm7* mRNA levels threefold in cells treated with rMO-CTL, whereas this effect was attenuated in cells pretreated with rMO-R1 **(Figure 2A)**. Consistent with these findings, rMO-R1 abolished leptin-induced demethylation at R1 but not R2 and R3 **(Figure 2B-D)**.

**Figure 2.**
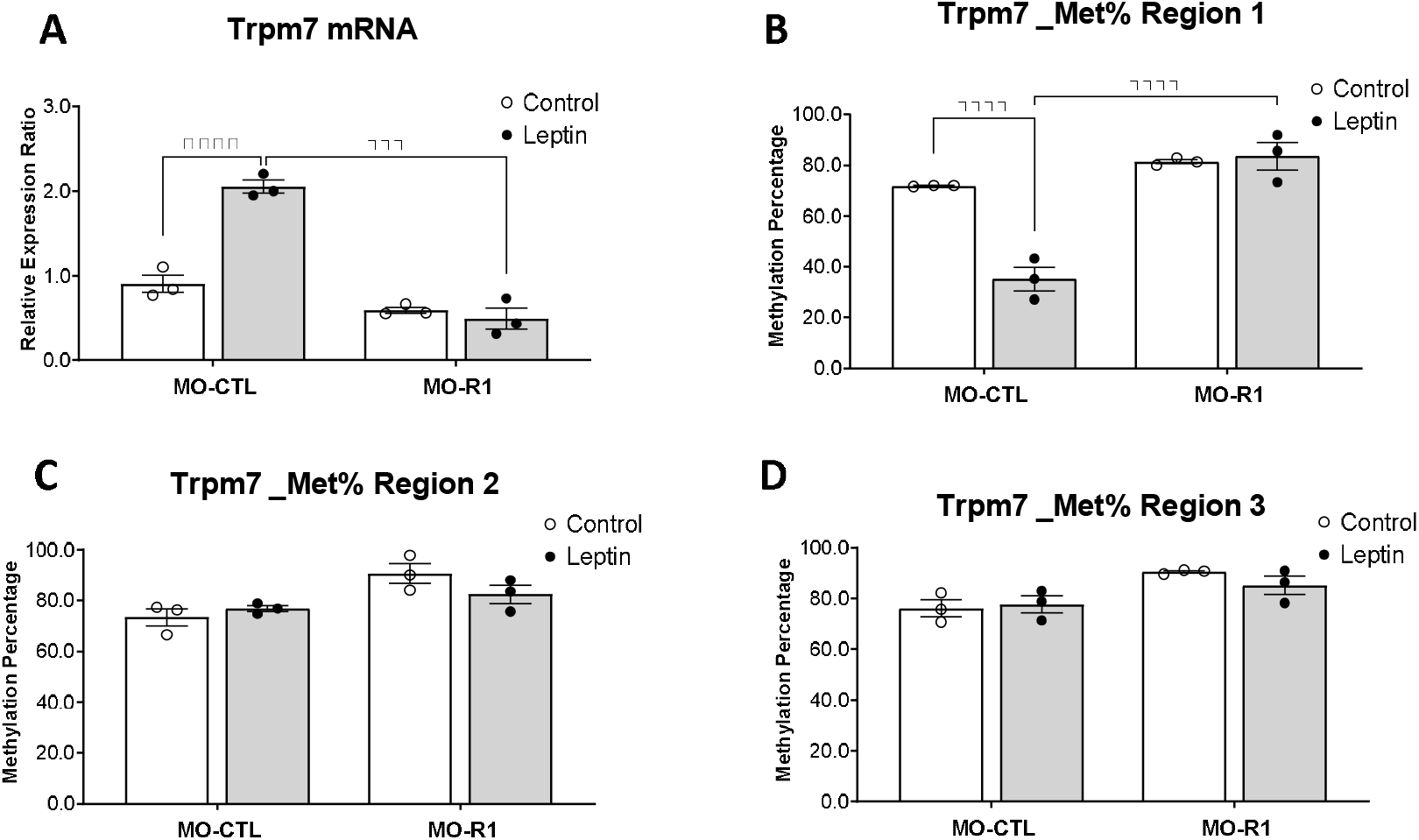
Methylated DNA oligonucleotides for Region 1 (R1) induced CpG site-specific DNA methylation and attenuated leptin-induced Trpm7 transcription in rat PC12 cells. PC12 cells were treated with methylated DNA oligonucleotides for control sequence (MO-CTL) or R1 (MO-R1) prior to 48-hour treatment of 1 ng/mL leptin. **(A)** Trpm7 mRNA expression and **(B-D)** region-specific DNA methylation were assessed using qPCR and Methylation-Specific PCR (MSPCR), respectively. All data are presented as mean ± SEM.

We next applied this strategy *in vivo* by administering methylated DNA oligonucleotides targeting R1 to the CBs of DIO mice. MO-R1 treatment increased DNA methylation at R1 but not at R2 or R3 **(Figure 3A)**, which correlated with reduced Trpm7 expression in the CB, but not in the medulla and hypothalamus tissues **(Figure 3B-D)**. Body weight, food intake, body temperature and plasma leptin levels did not differ between MO-R1- and MO-CTL-treated mice **(Table 1)**. These results indicate that MO-R1 specifically downregulates *Trpm7* expression in the CB *via* CpG site-specific promoter methylation.

**Figure 3.**
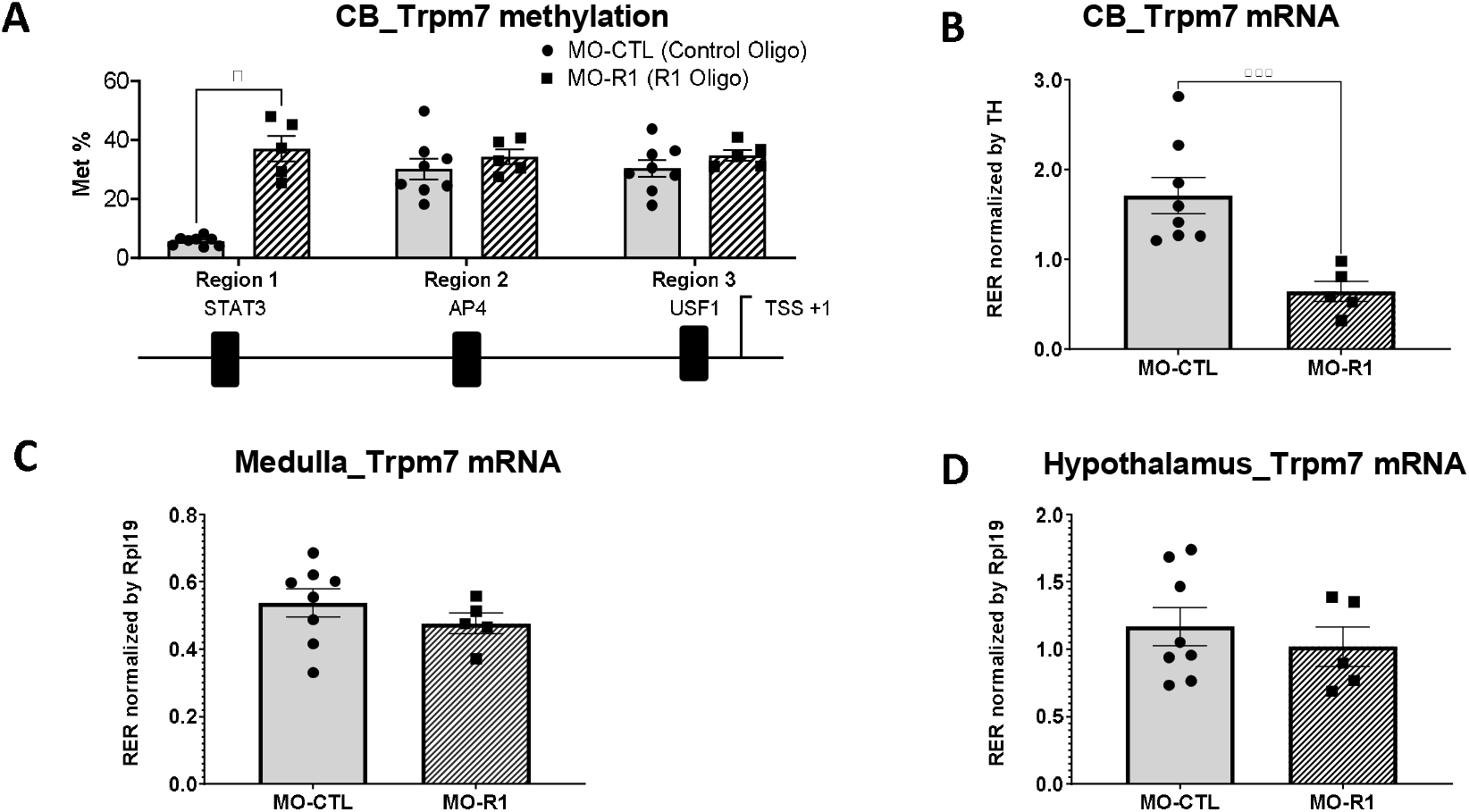
Methylated DNA oligonucleotides for Region 1 (R1) induced CpG site-specific DNA methylation and attenuated Trpm7 transcription in CB tissues of DIO mice. DIO mice were exposed to methylated DNA oligonucleotides for control sequence (MO-CTL) or R1 (MO-R1). **(A)** Region-specific DNA methylation and **(B)** mRNA levels of Trpm7 in CB tissues were assessed using Methylation-Specific PCR (MSPCR) and qPCR, respectively. **(C-D)** Trpm7 mRNA expression in medulla and hypothalamus tissues were also measured by qPCR. All data are presented as mean ± SEM. *TH*, tyrosine hydroxylase. Rpl19, ribosomal protein L19. Each dot represents an individual mouse.

**Table 1.** Basic characteristics of diet-induced obese (DIO) male mice before and after methylated oligonucleotides specific for the Trpm7 promoter Region 1 (MO-R1) or control (MO-CTL) oligos application to the carotid bifurcation area. There were no significant effects of MO-R1 or MO-CTL oligos applications.

|  | **MO-CTL** |  | **MO-R1** |  |
| --- | --- | --- | --- | --- |
| N | 13 |  | 11 |  |
| Time | Before surgery | After surgery | Before surgery | After surgery |
| Age (weeks) | 22.8 ± 0.3 | 25.8 ± 0.4 | 24 ± 0.1 | 26 ± 0.1 |
| Body Weight (g) | 46.3 ± 2.6 | 49.9 ± 2.2 | 49.2 ± 1.7 | 49.5 ± 1.8 |
| Rectal temperature (°C) | 38.5 ± 0.1 | 38.2 ± 0.1 | 38.4 ± 0.3 | 38.0 ± 0.4 |
| Food intake (g/day) | 3.2 ± 0.4 | 3.5 ± 0.2 | 3.2 ± 0.1 | 3.5 ± 0.1 |
| Plasma leptin (ng/ml) | Not tested | 54 ± 6.5 | Not tested | 40.4 ± 4.59 |

Next, we determined the effect of MO-R1 on CSN activity. Mice were treated with methylated DNA oligonucleotides two weeks prior to sacrifice with CSN isolation. In DIO mice treated with MO-CTL, leptin perfusion (50 nM) significantly increased CSN activity *ex vivo* under normoxic conditions and augmented the hypoxic chemoreflex (ΔCSN activity). The effect of leptin was abolished by a Trpm7 inhibitor FTY720 (**Figure 4A**). In contrast, MO-R1 treatment abolished the effects of leptin and FTY720 on CSN activity under both normoxic and hypoxic conditions (**Figure 4B**).

**Figure 4.**
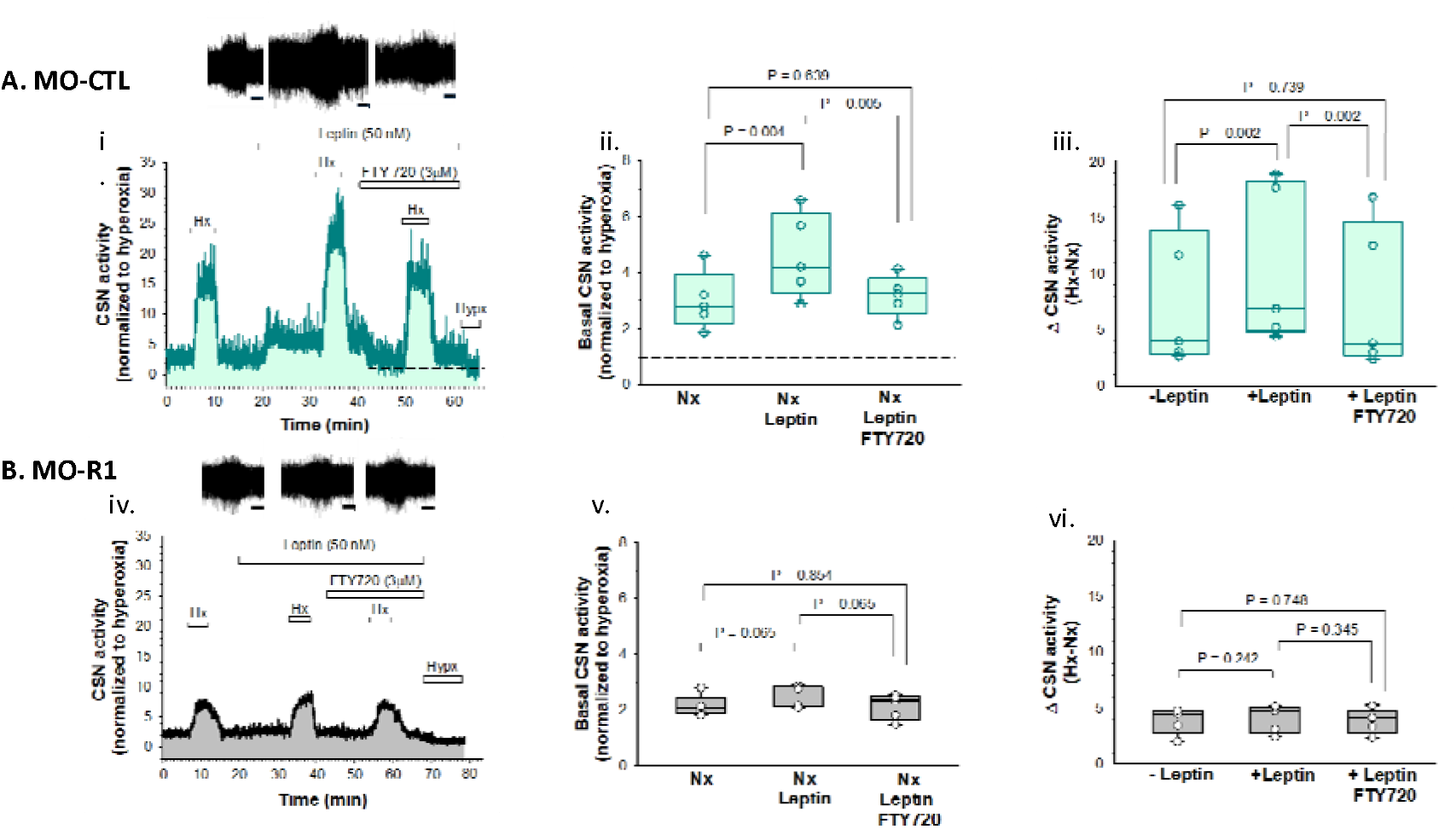
The effect of control methylated oligonucleotides (MO-CTL) and methylated oligonucleotides targeting the Region 1 of the Trpm7 promoter (MO-R1) on Carotid Sinus Nerve (CSN) activity in diet-induced obese mice. **(A**) (i) Representative integrated CSN activity trace in a CB preparation from an MO-CTL-treated mouse; showing the sequential effects of leptin and a TRPM7 blocker FTY720 during normoxia and hypoxia. Raw tracings throughout the time course (X-axis) are shown at the top; the scale bars represent 2 min. (ii) Summary data showing leptin induced significant increase in basal CSN activity (P = 0.004), which was abolished by Trpm7 receptor inhibitor FTY 720 (P = 0.005; N=5). (iii) Summary data showing CB chemosensitivity Hx (hypoxia) – Nx (normoxia), which was markedly enhanced by leptin (P = 0.002) and reversed by FTY 720 (P = 0.002; N=5). (**B)**. (iv) Representative integrated CSN activity in a preparation from an MO-R1 treated mouse showing the sequential effects of leptin and a TRPM7 blocker FTY720 during normoxia and hypoxia. Raw tracings throughout the time course (X-axis) are shown at the top; the scale bars represent 2 min. (v) Summary data of CSN activity under normoxia, which did not significantly increase by leptin (P = 0.065; N=5) and was unaffected by FTY 720 (P = 0.065; N = 5). (iv) Summary data showing CB chemosensitivity responses (Hx − Nx) remained unchanged across treatments (P = 0.451; N= 5). Values are expressed as box and whisker plots (median, 25-75% centiles and minimum and maximum values; mean ± S.D.). Data were statistically compared by one-way repeated measures ANOVA with Holm-Sidak post-hoc comparisons. P< 0.05 was considered significant.

MO-CTL treatment had no effect on blood pressure or heart rate (**Figures 5 and 6**). In contrast, compared to pretreatment levels, MO-R1 significantly decreased mean arterial pressure during the light phase when animals were predominantly asleep, from 104.7 ± 1.3 mm Hg to 99.3 ± 1.7 mm Hg (p = 0.001; **Figures 5 and 6**). The effect was present both for the light phase systolic pressure, which decreased from 130.4 ± 2.3 to 124.8 ± 1.9 mm Hg (p < 0.001), and diastolic pressure, which decreased from 91.8 ± 1.1 to 86.6 ± 1.7 mm Hg (p < 0.01). Heart rate decreased from 553 ± 15 to 523 ± 17 beats/min (p < 0.05). Smaller reductions in systolic pressure were observed during the active dark phase, whereas diastolic pressure and heart rate were unchanged. Overall, MO-R1 produced modest but statistically significant reductions in 24-hour mean and systolic arterial pressure.

**Figure 5.**
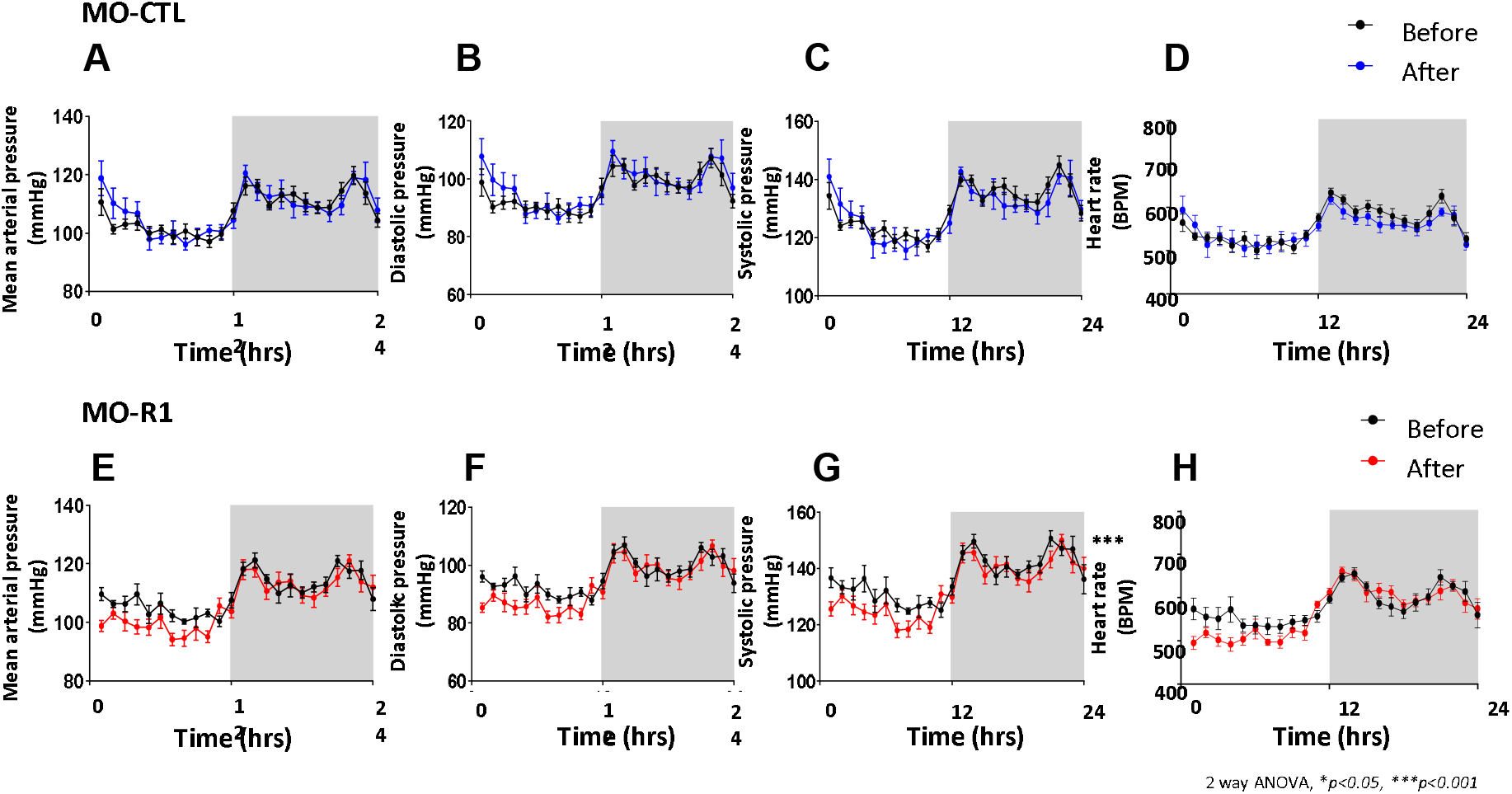
The effect of untargeted methylated oligonucleotides (MO-CTL, A-D) and methylated oligonucleotides targeting the Region 1 of the Trpm7 promoter (MO-R1, E-H) in the carotid bodies on the mean (A, E), systolic (B, F), and diastolic (C, G) arterial blood pressure and heart rate (D, H) profiles over 24 hours in diet-induced obese mice. Each dot represents average blood pressure values per group over 1 h interval in the control (MO-CTL, N = 8) and MO-R1 oligo (N = 6) treatments. The dark phase is shaded. *p < 0.05, ***, p < 0.001 between the before and after treatment in R1 oligo group. Repeated measures ANOVA.

**Figure 6.**
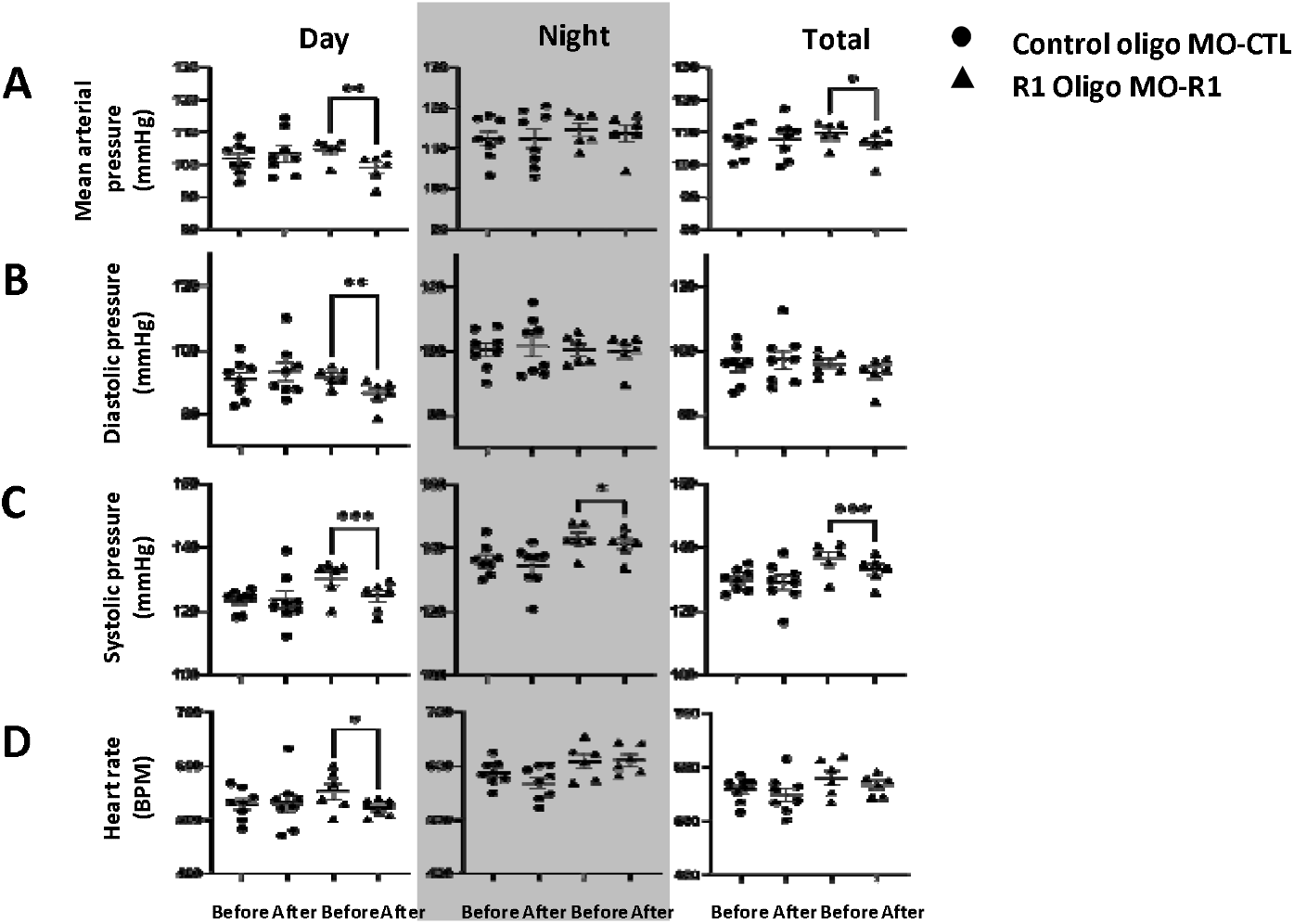
The effect of untargeted methylated oligonucleotides (MO-CTL, A-D, N = 8) and methylated oligonucleotides targeting the Region 1 of the Trpm7 promoter (MO-R1, E-H, N = 6) in the carotid bodies on mean (A), systolic (B), and diastolic (C) blood pressure and heart rate (D, HR) in individual mice. The dark phase is shaded. Each dot represents one animal. ** p < 0.01, respectively. Two-way ANOVA.

## Discussion

Obesity is a major cause of resistant hypertension, which has been attributed, at least in part, to high circulating levels of leptin (41, 57, 58). We previously reported that obesity-associated hyperleptinemia induces demethylation of the *Trpm7* promoter, which leads to upregulation of the *Trpm7* gene expression in the CB glomus cells, increased non-selective cation glomus cell current and CSN activity, and ultimately elevated blood pressure (19, 20). Using an *in vitro* model, we further showed that leptin induces region-specific demethylation of the *Trpm7* promoter, including a region encompassing the STAT3 binding site (R1) (48). Leptin signaling *via* LEPR^b^ is known to induce STAT3 phosphorylation (38, 39). The principal finding of the current study is that *in vivo* CB administration of methylated DNA oligonucleotides targeting the pSTAT3 binding region restored *Trpm7* promoter methylation, reduced *Trpm7* gene expression in the CB, abolished leptin-induced augmentation of CSN activity and lowered blood pressure in DIO mice.

Our findings provide mechanistic evidence that epigenetic modifications of CB genes can activate the CB-CSN-SNS axis and promote hypertension. The role of epigenetic regulation of CB in neural mechanisms of cardiorespiratory control has been explored previously. Rats exposed to intermittent hypoxia exhibit increased DNA methylation and repression of several antioxidant genes including superoxide dismutase 1 and 2, catalase, thioredoxin reductase 2, peroxiredoxin and glutathione peroxidase 2 in the CB and brainstem regions associated involved in CB reflexes (59, 60). Perinatal nicotine exposure induces hypertension in rat offspring through epigenetic upregulation of angiotensin signaling in CB (61). Obesity and comorbid OSA induce hypertension through CB-dependent mechanisms (19–21, 30, 62). We have previously identified an adipose-tissue produced hormone leptin as a key mediator of hypertension in obese rodents (19, 20, 47), and showed that leptin acts in the CB by upregulating the TRPM7 channel in type I glomus cells (19). We have also demonstrated *in vivo* that obesity induces demethylation of the *Trpm7* promoter (36). However, prior studies lacked direct mechanistic evidence that correction of epigenetic alterations can attenuate CB-CSN activity.

Targeted inhibition of transcription using short CpG-methylated DNA oligonucleotides, which induce locus-specific DNA methylation, was originally developed to silence the oncogene *Bcl-2 in vitro* (63). We previously demonstrated that methylated DNA oligonucleotides targeting *PDE4D* promoter reversed *PDE4D* upregulation and abnormal airway smooth muscle phenotypes in patients with asthma (54). Administration of a methylated oligonucleotide complementary to a specific *IGF2* promoter region increased survival in humanized mouse model of hepatocellular carcinoma (64). However, the targeted gene promoter methylation approach has not been previously tested *in vivo* in non-oncologic disease models. In the present study, methylated DNA oligonucleotides complementary to the STAT3 binding region of the *Trpm7* promoter (MO-R1), administered locally to the CB, reduced blood pressure in DIO mice. The hypotensive effect was most pronounced during the light phase, when mice are predominantly asleep, with mean arterial pressure decreasing by more than 5 mm Hg. Physiological nocturnal blood pressure dipping is an important protective mechanism against cardiovascular disease in humans (65). Loss of dipping is associated with increased sympathetic activity and is particularly prevalent in OSA (65–68). We previously demonstrated that DIO mice develop sleep disordered breathing resembling human OSA and obesity hypoventilation syndrome (37). In the present study, MO-R1 significantly reduced *Trpm7* expression in the CB, which we have shown improves sleep disordered breathing (36). Thus, the antihypertensive effect of MO-R1 may be mediated, at least in part, by improvement in sleep disordered breathing and attenuation of intermittent hypoxia (69).

*Ex vivo* measurements of CSN activity provided an additional mechanistic insight. Methylation of the pSTAT3 binding site within the *Trpm7* promoter prevented leptin-induced augmentation of CSN activity under both normoxic and hypoxic conditions, without affecting baseline CB activity or the hypoxic chemoreflex. Taken together with marked suppression of *Trpm7* expression by MO-R1 **(Fig. 4)**, these findings are consistent with our recent report showing that CB-specific *Trpm7* knockdown abolishes leptin-induced CSN activation in obese mice (70). Although multiple pathways may contribute to CB sensitization in obesity, including low-grade inflammation (71) and insulin resistance (21, 28), elevated circulating leptin appears to play a central role by inducing site-specific demethylation of the *Trpm7* promoter.

To our knowledge, this study provides the first evidence that targeted CpG site-specific promoter methylation can be used to lower blood pressure in obesity. These findings suggest a novel, potentially personalized epigenetic approach to treatment of hypertension, a highly prevalent condition that is frequently resistant to therapy. From a broader prospective, this strategy may be applicable to other non-oncologic diseases; however, several limitations should be acknowledged. First, MO-R1 was administered locally to the CB area, which presents a barrier to clinical translation. Although *Trpm7* is ubiquitously expressed and its systemic inhibition may confer additional benefits, including seizure suppression (72) and neuroprotection after stroke (73, 74), systemic delivery would likely require higher doses and potential toxicity must be carefully evaluated. Second, blood pressure and CSN activity were assessed two weeks after MO-R1 administration, and the duration of the therapeutic effect remains unknown. Third, we did not test methylated DNA oligonucleotides targeting other sites of the *Trpm7* promoter (R2 and R3), which have been shown to undergo demethylation in response to leptin in PC12 cells. Finally, the role of CB *Trpm7* CB inhibition and epigenetic modulation has not been examined in other models of hypertension, including spontaneously hypertensive rats, intermittent hypoxia-induced, or renovascular hypertension.

## Conclusions and Implications

Administration of methylated DNA oligonucleotides complementary to the pSTAT3 binding site of the *Trpm7* promoter to the carotid body reduced local *Trpm7* expression, abolished leptin-induced augmentation of CSN activity at normoxic and hypoxic conditions, and lowered blood pressure in DIO mice. These findings support epigenetic targeting of the *Trpm7* promoter as a potential therapeutic strategy for obesity-induced hypertension.

## Methods

### Ethical Approval

All experimental procedures were approved by The George Washington University School of Medicine Animal Care and Use Committee under protocol number A2023-025. All surgical manipulations involving carotid body (CB) microinjections were performed under aseptic conditions. Animals were anesthetized with 2% isoflurane for induction and maintained under 1.0–1.5% isoflurane (flow rate 2 L·min^−1^) throughout the surgical procedures. Buprenorphine-SR (3.25 mg·kg^−1^, subcutaneous) was administered 30–60 minutes prior to surgery for analgesia and re-administered 72 hours later if any signs of distress or pain were observed. All recovery procedures were carefully monitored to ensure animal welfare and compliance with institutional guidelines.

### Experimental Animals

A total of twenty-four male diet-induced obese (DIO) mice on the C57BL/6J background were purchased at 18 weeks of age from The Jackson Laboratory (Bar Harbor, ME, USA; stock #380050). DIO does not induce hypertension in female mice. Moreover, LEPR^b^ knockdown in CB interrupting the leptin-TRPM7 pathway does not decrease blood pressure in DIO female mice (47). Therefore, only DIO male mice were used in this study. Mice were fed a high-fat diet (Research Diets D12492, New Brunswick, NJ, USA; 5.21 kcal·g^−1^, 60% kcal from fat and 20% kcal from carbohydrates) until they reached 22–26 weeks of age, achieving an average body weight of 46–50 g. They were housed in the standard laboratory environment at 22^°^C, the 12:12-h light-dark cycle (8 am to 8 pm lights on) with free access to food and water.

### Chemicals and Reagents

The physiological buffer used for carotid body (CB) dissection and perfusion contained (in mM): 115 NaCl, 4 KCl, 24 NaHCO_3_, 1.25 NaH_2_PO_4_, 2 CaCl_2_, 10 D-glucose, and 12 sucrose (Sigma-Aldrich, St. Louis, MO, USA). Leptin (Cat. #498-OB; R&D Systems, Minneapolis, MN, USA), FTY720 (Cat. #SML0700; Sigma-Aldrich, St. Louis, MO, USA), and isoflurane (Item #502017; MWI Animal Health, AmerisourceBergen; Boise, ID, USA) were used in all experimental protocols.

### Assessment of Trpm7 gene expression and DNA methylation levels in mouse CB tissues

Total RNA from CB, medulla and hypothalamus of lean (C57BL/6J) or DIO mice was extracted using Trizol and PicoPure RNA isolation kit (ThermoFisher Scientific, MA). cDNA was generated using High-Capacity cDNA kit (ThermoFisher Scientific, MA) and subject for PCR with GoTaq Probe qPCR Mix (Promega, WI) and TaqMan Gene Expression Assays (ThermoFisher Scientific, MA) with the following conditions: 95°C for 3 minutes, followed by 45 cycles of 95°C for 30 seconds and 60°C for 1 minute. The mRNA level of *Trpm7* (Mm00457998_m1) was normalized to the levels of *Rpl19* (Mm02601633_g1). The 2-ΔΔCt method was used to calculate the relative expression level of the transcripts. qPCR results were present as mean values. For CB tissues, *Trpm7* mRNA level was normalized to the levels of tyrosine hydroxylase (TH, Mm00447557_m1). TH has been previously validated as a reference gene in glomus cells by Shin et al.(19) showing no significant changes in TH mRNA or protein levels in lean and DIO mice.

We previously employed bisulfite genomic sequencing and identified leptin induced region-specific DNA demethylation at rat *Trpm7* promoter (48). Using the mouse homolog of Trpm7 genomic sequence, we designed three sets of primers targeting specific methylated CpG sites at three regions within mouse *Trpm7* promoter. *In silico* analysis revealed putative transcription factor (TF) binding sites within these regions. Primer sequences were MSPCR_R1-F: 5’-TTGAAATTTTAGTTTTGGAGGATTTAGTACG -3’ for forward and MSPCR_R1-R: 5’-GCATCTATCTACGTTAATTATTAACGAT -3’ for reverse synthesis; MSPCR_R2-F: 5’-TGTATTTTTGATGGGAAAGGATAAC -3’ for forward and MSPCR_R2-R: 5’-GTAAAAAACGCGCTTACTACGAC -3’ for reverse synthesis; and MSPCR_R3-F: 5’-GTAGGTCGTAGTAAGCGCGT -3’ for forward and MSPCR_R3-R: 5’-GAAAAACGTCTCCGAAAACG -3’ for reverse synthesis. Genomic DNA was used for bisulfite treatment using the EZ DNA Methylation™ Kit (Zymo Research, CA). The bisulfite treated DNA was amplified with methylation specific primers using GoTaq® qPCR Master Mix (Promega, WI) and optimized PCR conditions (95°C for 3 minutes, 42 cycles of 95°C for 30 seconds, 59°C for 1 minute and 72°C for 2 minutes). Beta Actin was used as a reference gene to normalize the amount of bisulfite-treated DNA template using primers, MSPCR-Actin-F (5’AGT AAT AGT TGG AAA GTT AGA TTT-3’) and MSPCR-Actin-R (5’ ACC ACT AAA ACT CAC CCT ATA-3’). Fully methylated control DNA (Zymo Research, Irvine, CA) was used as a reference to calculate the percentage methylation of DNA samples.

### Assessment of rat *Trpm7* gene expression and DNA methylation levels in rat PC12 cells

PC12^LEPRb^ cells were cultured as described (48). Cells were transfected with non-targeting methylated control DNA (rMO-CTL) or DNA oligonucleotides (rMO-R1) specific for methylated region of *Trpm7* promoter (targeting CpG sites differentially demethylated by leptin as described in Yeung et al., RCMB 2021) prior to 48-hour leptin (1ng/mL). The phosphorothioate oligonucleotides were designed to replace the cytosines in CpG dinucleotides with methylated cytosine. Sequences for rMO-CTL and rMO-R1 are TGC/iMe-dC/CGGAA/iMe-dC/CGTGAGCACTT/iMe-dC/CTG and GATGATCTAGGG/iMe-dC/CGGTCTTC/iMe-dC/CGA respectively. DNA oligonucleotides were purchased from Integrated DNA Technologies (Coralville, IA). DNA and DNA from PC12 cells were isolated using the AllPrep DNA/RNA/miRNA Universal Kit (Qiagen). The 2-ΔΔCt method was used to calculate the relative expression ratio (RER) of Trpm7 transcript that normalized by housekeeping gene Rpl19. Genomic DNA was used for bisulfite treatment using the EZ DNA Methylation™ Kit (Zymo Research, CA). The bisulfite treated DNA was subject for DNA methylation assays to assess the region-specific DNA methylation changes within rat *Trpm7* promoter as previously described. The primers used for gene expression and DNA methylation studies were reported by Yeung et al (48).

### Methylated DNA oligonucleotides application to the carotid body (CB) area of DIO mice

After validating the effectiveness of methylated DNA oligonucleotides in targeting leptin-induced CpG site demethylation in PC12 cells, we applied a similar approach to target a demethylated CpG site within mouse *Trpm7* promoter to reduce *Trpm7* transcription. Animals were premedicated with buprenorphine XR (Ethiqa) at 3.25 mg/kg. Methylated DNA oligonucleotides for R1 (MO-R1) (n = 13 mice) or non-targeting control (MO-CTL) (n = 11 mice) were applied in DIO mice under 2 - 3 % isoflurane anesthesia as previously described(75). Briefly, the bifurcations of the common carotid arteries were exposed bilaterally. MO-R1 (5’-A/iMe-dC/CGGTCTTCTGAGGACACTGCA-3’) or MO-CTL (5’-TGC/iMe-dC/CGGAACGTGAGCACTTCTG-3’) from Integrated DNA Technologies (Coralville, IA) were suspended in 5 µl of mixture by in vivo-jetPEI from Polyplus (New York, NY). After the oligo suspension was congealed (2-3 min), the incision was closed. Mice recovered for 14 days and BP was measured in 24 mice for 24 hs prior to sacrifice (n = 8 in the MO-CTL group and n = 6 in the MO-R1 group), whereas 10 mice (n= 5 per group) have been used for the CB-CSN preparation. Upon completion of the measurements (14 days after transfection), mice were sacrifice; CB, the brain medulla and hypothalamus were harvested in liquid nitrogen and then stored at -80^°^C. Gene expression and DNA methylation changes were assessed as abovementioned.

### CSN activity Measurements *Ex vivo*

#### Ex Vivo Carotid Body–Carotid Sinus Nerve (CB–CSN) Preparation

Ten DIO mice have been used in the CB-CSN experiments. Heavily anesthetized mice were decapitated at the lower cervical level, and the head was immediately transferred to ice-cold, oxygenated artificial cerebrospinal fluid (aCSF; equilibrated with 95% O_2_ and 5% CO_2_). The carotid bifurcation, including the CB, carotid sinus nerve (CSN), and superior cervical ganglion, was rapidly isolated en bloc as described previously (Roy et al., 2012). Dissections were performed in chilled aCSF (115 mM NaCl, 4 mM KCl, 24 mM NaHCO_3_, 2 mM CaCl_2_, 1.25 mM NaH_2_PO_4_, 1 mM MgSO_4_, 10 mM glucose, and 12 mM sucrose), continuously bubbled with 95% O_2_/5% CO_2_ to maintain pH ≈ 7.4. Following dissection, the carotid bifurcation was transferred to a custom-built recording chamber equipped with a water-jacketed heating system. The common carotid artery was cannulated for luminal perfusion with aCSF equilibrated to 100 mmHg PO_2_ and 35 mmHg PCO_2_ (balance N_2_). Perfusion was maintained using a peristaltic pump at a flow rate of 7 mL·min^−1^, yielding a stable intraluminal pressure of 90–100 mmHg at the cannula tip. All branches of the carotid bifurcation (occipital, internal, and external carotid arteries) were ligated, and small incisions (except occipital artery) were made to facilitate perfusate exit. Before entering the preparation, the perfusate was passed through a bubble trap and a heat exchanger. The perfusion temperature was continuously monitored in the recording chamber and maintained at 37 ± 0.5°C. The effluent from the chamber was recirculated.

#### Carotid Sinus Nerve (CSN) Recording

Chemosensory activity was recorded using a suction electrode; signals were rectified and integrated with a 200 ms time constant. A reference electrode was positioned near the carotid bifurcation. Neural activity was amplified and filtered (0.3–1 kHz) using a differential AC amplifier (Model 1700, A-M Systems). Signals were rectified, integrated (200 ms time constant), and digitized using a Digidata 1322A interface (Axon Instruments) with AxoScope 9.0 software for acquisition and analysis. CB-CSN preparations that displayed spontaneous action potential under baseline condition were considered viable, and recordings were obtained from one preparation per animal. Each preparation was stabilized for at least 30 minutes before initiating experimental protocols. MO-R1 and MO-CTL treated animals were subjected to identical experimental conditions and pharmacological interventions and the investigator (Dr. Roy) has been blinded to the treatment group.

#### Experimental Protocol

The experimental sequence for both groups consisted of the following steps:

i. 5 min normoxia (Nx; PO_2_ = 100 Torr) to establish baseline chemosensory activity.
ii. 5 min hypoxia (Hx; PO_2_ = 60 Torr).
iii. 5 min Nx recovery.
iv. 10 min Nx perfusion with leptin (50 nM in PBS).
v. 5 min Hx in presence of leptin.
vi. 5 min Nx in presence of leptin.
vii. 10 min Nx perfusion with FTY720 (3 μM in PBS) plus leptin.
viii. 5 min Hx in presence of FTY720 plus leptin.
ix. 5 min Nx recovery in presence of FTY720 plus leptin.
x. Final hyperoxic challenge (Hypx; PO_2_ = 200 Torr)

All recordings were performed continuously throughout the protocol. Each step was separated by adequate stabilization to ensure reproducibility and minimize drift in baseline activity.

### Cardiovascular telemetry

PA-C10 telemeters (Data Sciences, St. Paul, MN) were placed in the left femoral arteries under 2-3 % isoflurane as previously described(19). After 7-10 days recovery the baseline recording was performed. Signals were captured using PowerLabs 16/35 interfaced with LabChart Pro software from ADInstruments (Colorado Springs, CO).

### Biochemical assays

Terminal blood draws by cardiac puncture and tissue harvesting were performed under 2 - 3 % isoflurane anesthesia. The plasma was collected at the time of sacrifice and plasma leptin level were measured ELISA kits from Millipore (Billerica, MA) and compared between CON and R1 oligo applied mice.

### Statistical Analysis

Statistical analyses were structured to test a priori hypothesis that knockdown of demethylation of Trpm7 in the CB with R1 oligo will decrease blood pressure and carotid sinus nerve (CSN) activity.

CB neural recording and telemetry data were analyzed offline using LabChart 7 Pro (ADI Instruments).

#### CB-CSN responses

Raw CSN activity was divided into 1 s time bins, and the activity was expressed as integrated neural discharge and normalized to the hyperoxia. The final 1 min of CSN activity (the steady-state response) during each condition was averaged and used for statistical analysis. Differences in CSN activity between the experimental conditions were analyzed using one-way repeated-measures ANOVA method, followed by Holm–Sidak *post hoc* correction. Data were plotted using SigmaPlot version 15.0 (Systat Software San Jose, CA, USA) as box-and-whisker plots, with whiskers showing the minimal and maximal values, boxes outlining the second and third quartiles and the line showing the median value. In the text, results are presented as mean ± SD. *P <* 0.05 was considered statistically significant.

#### In vivo parameters including telemetry

Statistical significance for comparisons between MO-R1 and MO-CTL control groups before and after interventions was determined by two-way ANOVA with Bonferroni post-hoc correction for multiple comparisons and Tukey’s post hoc test. Other between group comparisons were analyzed with the Mann-Whitney test. The GraphPad Prism software has been used. A p-value of <0.05 was considered significant.

## References

1. Ogden CL, et al. Trends in Obesity Prevalence by Race and Hispanic Origin—1999-2000 to 2017-2018. JAMA. 2020;324(12):1208–1210.

2. Friedman AN, Ogden CL, Hales CM. Prevalence of Obesity and CKD Among Adults in the United States, 2017-2020. Kidney Medicine. 2023;5(1):100568.

3. Kotsis V, et al. Mechanisms of obesity-induced hypertension. Hypertension Research. 2010;33(5):386–393.

4. Rahmouni K, et al. Obesity-associated hypertension: new insights into mechanisms. Hypertension. 2005;45(1524-4563 (Electronic)):9–14.

5. Bramlage P, et al. Hypertension in overweight and obese primary care patients is highly prevalent and poorly controlled. AmJHypertens. 2004;17(0895-7061 (Linking)):904–910.

6. Franks PW, et al. Childhood obesity, other cardiovascular risk factors, and premature death. NEnglJMed. 2010;362(1533-4406 (Electronic)):485–493.

7. Isomaa B, et al. Cardiovascular Morbidity and Mortality Associated With the Metabolic Syndrome. Diabetes Care. 2001;24(4):683.

8. Grundy SM, et al. Definition of metabolic syndrome: Report of the National Heart, Lung, and Blood Institute/American Heart Association conference on scientific issues related to definition. Circulation. 2004;109(3):433–438.

9. O’Connor GT, et al. Prospective study of sleep-disordered breathing and hypertension: the Sleep Heart Health Study. American Journal of Respiratory & Critical Care Medicine. 2009;179(12):1159–1164.

10. Narkiewicz K, et al. Sympathetic activity in obese subjects with and without obstructive sleep apnea. Circulation. 1998;98(8):772–776.

11. Alberti KG, et al. Harmonizing the metabolic syndrome: a joint interim statement of the International Diabetes Federation Task Force on Epidemiology and Prevention; National Heart, Lung, and Blood Institute; American Heart Association; World Heart Federation; International Atherosclerosis Society; and International Association for the Study of Obesity. Circulation. 2009;120(1524-4539 (Electronic)):1640–1645.

12. Carey Robert M., et al. Prevalence of Apparent Treatment-Resistant Hypertension in the United States. Hypertension. 2019;73(2):424–431.

13. Young T, Skatrud J, Peppard PE. Risk factors for obstructive sleep apnea in adults. JAMA. 2004;291(16):2013–6.

14. Young T, Peppard PE, Gottlieb DJ. Epidemiology of Obstructive Sleep Apnea. American Journal of Respiratory & Critical Care Medicine. 2002;165:1217–1239.

15. Peppard PE, et al. Prospective study of the association between sleep-disordered breathing and hypertension. NEJM. 2000;342(19):1378–1384.

16. Tufik S, et al. Obstructive sleep apnea syndrome in the Sao Paulo Epidemiologic Sleep Study. Sleep Medicine. 2010;11(5):441–446.

17. Heinzer R, et al. Prevalence of sleep-disordered breathing in the general population: the HypnoLaus study. The Lancet Respiratory medicine. 2015;3(4):310–318.

18. Nieto FJ, et al. Association of sleep-disordered breathing, sleep apnea, and hypertension in a large community-based study: Sleep Heart Health Study. JAMA. 2000;283(14):1829–1836.

19. Shin M-K, et al. Leptin Induces Hypertension Acting on Transient Receptor Potential Melastatin 7 Channel in the Carotid Body. Circulation Research. 2019;125(11):989–1002.

20. Shin M-K, et al. Pharmacological and Genetic Blockade of Trpm7 in the Carotid Body Treats Obesity-Induced Hypertension. Hypertension. 2021;78(1):104–114.

21. Shin M-K, et al. Carotid body denervation improves hyperglycemia in obese mice. Journal of Applied Physiology. 2024;136(2):233–243.

22. Paton JFR, et al. The Carotid Body as a Therapeutic Target for the Treatment of Sympathetically Mediated Diseases. Hypertension. 2013;61(1):5–13.

23. Accorsi-Mendonça D, et al. Enhanced Firing in NTS Induced by Short-Term Sustained Hypoxia Is Modulated by Glia-Neuron Interaction. J Neurosci. 2015;35(17):6903.

24. Nurse CA, Piskuric NA. Signal processing at mammalian carotid body chemoreceptors. SeminCell DevBiol. 2013;24(1096-3634 (Electronic)):22–30.

25. Thakkar P, et al. Carotid body: an emerging target for cardiometabolic co-morbidities. Experimental Physiology. 2023;108(5):661–671.

26. Pijacka W, et al. Purinergic receptors in the carotid body as a new drug target for controlling hypertension. NatMed. 2016;22(1546–170X (Electronic)):1151–1159.

27. Abdala AP, et al. Hypertension is critically dependent on the carotid body input in the spontaneously hypertensive rat. JPhysiol. 2012;590(1469-7793 (Electronic)):4269–4277.

28. Ribeiro MJ, et al. Carotid Body Denervation Prevents the Development of Insulin Resistance and Hypertension Induced by Hypercaloric Diets. Diabetes. 2013;62(8):2905.

29. Shin MK, et al. Carotid body denervation prevents fasting hyperglycemia during chronic intermittent hypoxia. Journal of applied physiology (Bethesda, MdrI: 1985). 2014;117(7):765–776.

30. Fletcher EC, et al. Carotid chemoreceptors, systemic blood pressure, and chronic episodic hypoxia mimicking sleep apnea. J Appl Physiol (1985). 1992;72(5):1978–1984.

31. Niewinski P, et al. Carotid body resection for sympathetic modulation in systolic heart failure: results from first-in-man study. European Journal of Heart Failure. 2017;19(3):391–400.

32. Fudim M, et al. Effects of carotid body tumor resection on the blood pressure of essential hypertensive patients. Journal of the American Society of Hypertension. 2015;9(6):435–442.

33. Narkiewicz K, et al. Unilateral Carotid Body Resection in Resistant Hypertension: A Safety and Feasibility Trial. Jacc Basic to Translational Science. 2016;1(5):313–324.

34. Considine RV, et al. Serum immunoreactive-leptin concentrations in normal-weight and obese humans. New England Journal of Medicine. 1996;334(5):292–295.

35. Maffei M, et al. Leptin levels in human and rodent: measurement of plasma leptin and ob RNA in obese and weight-reduced subjects. NatMed. 1995;1(11):1155–1161.

36. Kim LJ, et al. TRPM7 channels regulate breathing during sleep in obesity by acting peripherally in the carotid bodies. The Journal of Physiology. 2022;600(23):5145–5162.

37. Fleury Curado T, et al. Sleep-disordered breathing in C57BL/6J mice with diet-induced obesity. Sleep. 2018;41(8):zsy089.

38. Bjorbaek C, et al. Divergent signaling capacities of the long and short isoforms of the leptin receptor. Journal of Biological Chemistry. 1997;272(51):32686–32695.

39. Bates SH, et al. STAT3 signalling is required for leptin regulation of energy balance but not reproduction. Nature. 2003;421(6925):856–859.

40. Rahmouni K, et al. Role of selective leptin resistance in diet-induced obesity hypertension. Diabetes. 2005;54(0012-1797 (Print)):2012–2018.

41. Asferg C, et al. Leptin, not adiponectin, predicts hypertension in the Copenhagen City Heart Study. AmJHypertens. 2010;23(1941-7225 (Electronic)):327–333.

42. Shankar A, Xiao J. Positive relationship between plasma leptin level and hypertension. Hypertension. 2010;56(1524-4563 (Electronic)):623–628.

43. Hall JE, et al. Obesity-induced hypertension: role of sympathetic nervous system, leptin, and melanocortins. JBiolChem. 2010;285(1083–351X (Electronic)):17271–17276.

44. Hall JE, et al. Obesity, kidney dysfunction, and inflammation: interactions in hypertension. Cardiovascular Research. 2021;117(8):1859–1876.

45. Caballero-Eraso C, et al. Leptin acts in the carotid bodies to increase minute ventilation during wakefulness and sleep and augment the hypoxic ventilatory response. The Journal of Physiology. 2019;597(1):151–172.

46. Kim LJ, et al. Leptin Receptor Blockade Attenuates Hypertension, but Does Not Affect Ventilatory Response to Hypoxia in a Model of Polygenic Obesity. Frontiers in Physiology. 2021;12:1042.

47. Shin M-K, et al. Leptin receptor downregulation in the carotid body treats obesity-induced hypertension. Journal of Neurophysiology. 2025;133(3):892–903.

48. Yeung BH, et al. Leptin Induces Epigenetic Regulation of Transient Receptor Potential Melastatin 7 in Rat Adrenal Pheochromocytoma Cells. Am J Respir Cell Mol Biol. 2021;65(2):214–221.

49. Ahuja N, Sharma AR, Baylin SB. Epigenetic Therapeutics: A New Weapon in the War Against Cancer. Annual Review of Medicine. 2016;67(Volume 67, 2016):73–89.

50. Majchrzak-Celińska A, Warych A, Szoszkiewicz M. Novel Approaches to Epigenetic Therapies: From Drug Combinations to Epigenetic Editing. Genes. 2021;12(2). 10.3390/genes12020208.

51. Talukdar PD, Chatterji U. Transcriptional co-activators: emerging roles in signaling pathways and potential therapeutic targets for diseases. Signal Transduction and Targeted Therapy. 2023;8(1):427.

52. Yao X, et al. A methylated oligonucleotide inhibits IGF2 expression and enhances survival in a model of hepatocellular carcinoma. J Clin Invest. 2003;111(2):265–273.

53. Lei Y, et al. Targeted DNA methylation in vivo using an engineered dCas9-MQ1 fusion protein. Nature Communications. 2017;8(1):16026.

54. Lin AHY, et al. Aberrant DNA Methylation of Phosphodiestarase 4D Alters Airway Smooth Muscle Cell Phenotypes. Am J Respir Cell Mol Biol. 2016;54(2):241–249.

55. Klose RJ, Bird AP. Genomic DNA methylation: the mark and its mediators. Trends in Biochemical Sciences. 2006;31(2):89–97.

56. Jones PA. Functions of DNA methylation: islands, start sites, gene bodies and beyond. Nature Reviews Genetics. 2012;13(7):484–492.

57. Bell BB, Rahmouni K. Leptin as a Mediator of Obesity-induced Hypertension. Curr Obes Rep. 2016;5(4):397–404.

58. Kshatriya S, et al. Obesity hypertension: the regulatory role of leptin. Int J Hypertens. 2011;2011:270624–270624.

59. Nanduri J, et al. Epigenetic regulation of redox state mediates persistent cardiorespiratory abnormalities after long-term intermittent hypoxia. J Physiol (Lond). [published online ahead of print: August 9, 2016]. 10.1113/JP272346.

60. Nanduri J, Semenza GL, Prabhakar NR. Epigenetic changes by DNA methylation in chronic and intermittent hypoxia. American Journal of Physiology-Lung Cellular and Molecular Physiology. 2017;313(6):L1096–L1100.

61. Zhu F, et al. Epigenetic Upregulation of Carotid Body Angiotensin Signaling Increases Blood Pressure. Hypertension. 2025;82(2):293–305.

62. Prabhakar NR. Carotid body chemoreflex: a driver of autonomic abnormalities in sleep apnoea. Exp Physiol. 2016;101(8):975–985.

63. Hoffman AR, Hu JF. Directing DNA Methylation to Inhibit Gene Expression. Cellular and Molecular Neurobiology. 2006;26(4):423–436.

64. Yao X, et al. A methylated oligonucleotide inhibits IGF2 expression and enhances survival in a model of hepatocellular carcinoma. J Clin Invest. 2003;111(2):265–273.

65. Parati G, et al. Nocturnal blood pressure: pathophysiology, measurement and clinical implications. Position paper of the European Society of Hypertension. Journal of Hypertension. 2025;43(8). https://journals.lww.com/jhypertension/fulltext/2025/08000/nocturnal_blood_pressure__pathophysiology,.4.aspx.

66. Parati G, et al. Obstructive sleep apnea syndrome as a cause of resistant hypertension. Hypertension Research. 2014;37:601.

67. Elsanan MAHA, et al. Cardiovascular risks in non-dipper OSA patients: insights from ABPM, echocardiography, and Holter monitoring. Sleep and Breathing. 2025;29(3):193.

68. Genta-Pereira DC, et al. Nondipping Blood Pressure Patterns Predict Obstructive Sleep Apnea in Patients Undergoing Ambulatory Blood Pressure Monitoring. Hypertension. 2018;72(4):979–985.

69. Berger S, et al. Intranasal Leptin Relieves Sleep-disordered Breathing in Mice with Diet-induced Obesity. Am J Respir Crit Care Med. 2018;199(6):773–783.

70. Roy A, et al. Diet-induced obesity in mice increases carotid body chemosensitivity via the leptin-TRPM7 pathway. The Journal of Physiology. 2025;603(17):4887–4905.

71. Iturriaga R, Del Rio R, Alcayaga J. Carotid Body Inflammation: Role in Hypoxia and in the Anti-inflammatory Reflex. Physiology. 2022;37(3):128–140.

72. Khalil A, et al. Drugs acting at TRPM7 channels inhibit seizure-like activity. Epilepsia Open. 2023;8(3):1169–1174.

73. Chen W, et al. TRPM7 inhibitor carvacrol protects brain from neonatal hypoxic-ischemic injury. Molecular Brain. 2015;8(1):11.

74. Turlova E, et al. TRPM7 Mediates Neuronal Cell Death Upstream of Calcium/Calmodulin-Dependent Protein Kinase II and Calcineurin Mechanism in Neonatal Hypoxic-Ischemic Brain Injury. Translational Stroke Research. 2021;12(1):164–184.

75. Shin M-K, et al. Experimental Approach to Examine Leptin Signaling in the Carotid Bodies and its Effects on Control of Breathing. JoVE. 2019;(152):60298.

